# Breast cancer classification based on proteotypes obtained by SWATH mass spectrometry

**DOI:** 10.1101/583443

**Authors:** Pavel Bouchal, Olga T. Schubert, Jakub Faktor, Lenka Capkova, Hana Imrichova, Karolina Zoufalova, Vendula Paralova, Roman Hrstka, Yansheng Liu, H. Alexander Ebhardt, Eva Budinska, Rudolf Nenutil, Ruedi Aebersold

## Abstract

Accurate breast cancer classification is vital for patient management decisions, and better tumour classification is expected to enable more precise and eventually personalized treatment to improve patient outcomes. Here, we present a novel quantitative proteotyping approach based on SWATH mass spectrometry and establish key proteins for breast tumour classification derived from proteotype data. The study was based on 96 tissue samples representing five breast cancer subtypes according to conventional classification. Correlation of SWATH proteotype patterns indicated groups that largely recapitulate these subtypes. However, the proteotype-based classification also revealed varying degrees of heterogeneity within the conventional subtypes, with triple negative tumours being the most heterogeneous. Proteins that contributed most strongly to the proteotype-based classification include INPP4B, CDK1, and ERBB2, which are associated with oestrogen receptor status, tumour grade, and HER2 status, respectively. While these three key proteins exhibited high levels of correlation between protein and transcript levels (R>0.67), general correlation did not exceed R=0.29, indicating the value of protein-level measurements of biomarkers and disease-regulated genes. Overall, our data shows how large-scale protein-level measurements by next-generation proteomics can lead to improved patient stratification for precision medicine.

## Introduction

Despite the progress achieved in early cancer diagnosis and therapy, many patients develop fatal disease. This also applies to breast cancer, even though it is one of the best characterized malignant diseases. Breast cancer is currently classified into five intrinsic subtypes, typically using immunohistological markers (oestrogen receptor (ER), progesterone receptor (PR), *HER2* gene and/or ERBB2 protein status), tumour grade and/or proliferation. We will refer to these subtypes as “conventional subtypes”; they have been defined as follows: luminal A (ER^+^, HER2^−^, low proliferation), luminal B HER2^−^ (ER^+^, HER2^−^, high proliferation), luminal B HER2^+^ (ER^+^, HER2^+^, high proliferation), HER2 enriched (ER^−^, HER2^+^, high proliferation), and triple negative (ER^−^, PR^−^, HER2^−^, high proliferation) (Brouckaert et al., 2013; Lam et al., 2014; Parise and Caggiano, 2014). This classification guides decisions for the adjuvant therapy which, however, fails in a substantial proportion of cases due to cancer recurrence, therapy resistance, and/or metastasis (Parise and Caggiano, 2014). The development of advanced, generalized disease despite the therapy guided by the tumour classification into the subtypes described above indicates that the current classification scheme may not fully capture the genetic and molecular status of the cancer and that a refined classification system might better predict which patient groups respond best to the range of available therapies.

Nowadays, the search for better tumour classifiers significantly concentrates on the application of omics approaches, which are able to analyse thousands of genes, gene transcripts or proteins in a single experiment. The biochemical effector molecules in cells are proteins and their direct measurement is, therefore, in principle preferable over the inference of protein quantities from transcript measurements (expression arrays, RNA sequencing). However, the commonly used proteomic approaches based on mass spectrometry analysis in data-dependent acquisition (DDA) mode are often hampered by limited consistency and quantitative accuracy and are therefore less suitable for application to clinical cohorts of significant size. In contrast, targeted proteomic technologies overcome some of these limitations and provide improved quantification precision and reproducibility (Pernikarova and Bouchal, 2015). Kennedy and colleagues (Kennedy et al., 2014) recently demonstrated the ability of the targeted proteomic technique selected/multiple reaction monitoring (S/MRM) to quantify 319 breast cancer-associated proteins with high interlaboratory reproducibility. The data discriminated basal vs. luminal breast cancer phenotypes and largely correlated with oestrogen receptor levels in 30 cell lines.

To increase the number of proteins reproducibly quantified across samples, in the present study we use a highly multiplexed mode of targeted proteomics, Sequential Windowed Acquisition of All Theoretical Fragment Ion Spectra-Mass Spectrometry (SWATH-MS), a next-generation proteomics approach developed by Gillet and colleagues (Gillet et al., 2012). For the targeted analysis of the acquired data we built a comprehensive breast cancer-specific SWATH assay library. We applied the SWATH-MS technique to obtain digital proteome maps (or “proteotypes”) for a set of 96 breast tumour lysates (Data file S1) and classified them into five proteotype-based subtypes using a conditional reference tree algorithm (Hothorn et al., 2006). The algorithm found three key proteins that are highly effective for group separation; the agreement between our proteotype-based subtypes and the conventional subtypes is 84 %. The triple negative subtype showed the highest degree of heterogeneity of protein expression. In addition to allowing a more refined classification of breast cancer subtypes, the obtained SWATH-MS data allowed us to compare protein and transcript levels of over 2,700 genes. While the correlation of protein and transcript levels was low for most differentially expressed genes, it was strong for the three classifying proteins. This study is the first application of the SWATH-MS technique towards the generation of large-scale quantitative proteomics profiles of breast cancer tissues and confirms the potential of SWATH-MS to generate high-quality, information-rich data for improved tumour classification. Discrepancies between the classical tumour subtypes and our proteotype-based subtypes potentially indicates patients that would benefit of different treatment strategies.

## Results

### Generation of an assay library for quantifying breast cancer-associated proteins by SWATH-MS

To extract quantitative protein information from SWATH-MS datasets acquired from breast cancer patient tissue samples in a targeted manner, we generated an extensive spectral library based on samples of all classical breast cancer subtypes described above and fractionated pools thereof. From this spectral library we obtained reference spectra for 28,233 proteotypic peptides (FDR<0.01), representing 4,443 proteins (Data file S2A). This spectral library was used in the following to quantify proteins in breast cancer tissue lysates using the SWATH-MS approach. The assay library covers many key proteins involved in cancer-related pathways and molecular functions such as the cell cycle/p53, TGF-β, JAK-STAT, PI3K-AKT, EGFR, and Wnt pathways, as well as adherent junctions, ECM-receptor interactions, and apoptosis (Data file S2B). This breast cancer SWATH assay library has been made available through the SWATHAtlas database (www.SWATHAtlas.org) as a public resource to support further basic and applied breast cancer research.

### Generation of a SWATH-MS data matrix consisting of 2,842 consistently quantified proteins across 96 patient samples

We analysed the proteome of 96 breast cancer tumour tissues by SWATH-MS. Each tumour tissue was previously classified by a pathologist into one of the five conventional breast cancer subtypes (defined by ER, PR, HER2 status, and tumour grade) and according to their lymph node status. In addition to the analysis of individual breast cancer samples, we also analysed pooled samples of each of the five subtypes. For each subtype, lymph node negative and lymph node positive samples were pooled separately, generating ten sample pools in total. Using the SWATH assay library described above, we were able to extract quantitative data for 27,515 peptides and their modified variants representing 2,842 proteins across all individual samples. These 2,842 consistently quantified proteins cover the majority of molecular processes known to be involved in breast cancer (Fig. S1).

### Comparison of proteotype-based subtypes and conventional subtypes of breast cancer

Using the thus generated proteotypes for 96 samples we first asked to what extent tumour classification based on proteotypes correlated with the conventional subtype classification. We performed unsupervised hierarchical clustering on the proteotypes of the pooled samples. Fig. 1A shows that pools of lymph node positive and negative samples of each subtype clustered closely together, indicating high reproducibility of our measurements. Moreover, clustering revealed proteotype similarity between less aggressive luminal A and luminal B subtypes, whereas the more aggressive HER2 and triple negative subtypes formed a separate cluster. The luminal B HER2^+^ group was more similar to the cluster with high aggressiveness, in agreement with its worse therapy response (Fig. 1A).

**Fig. 1.**
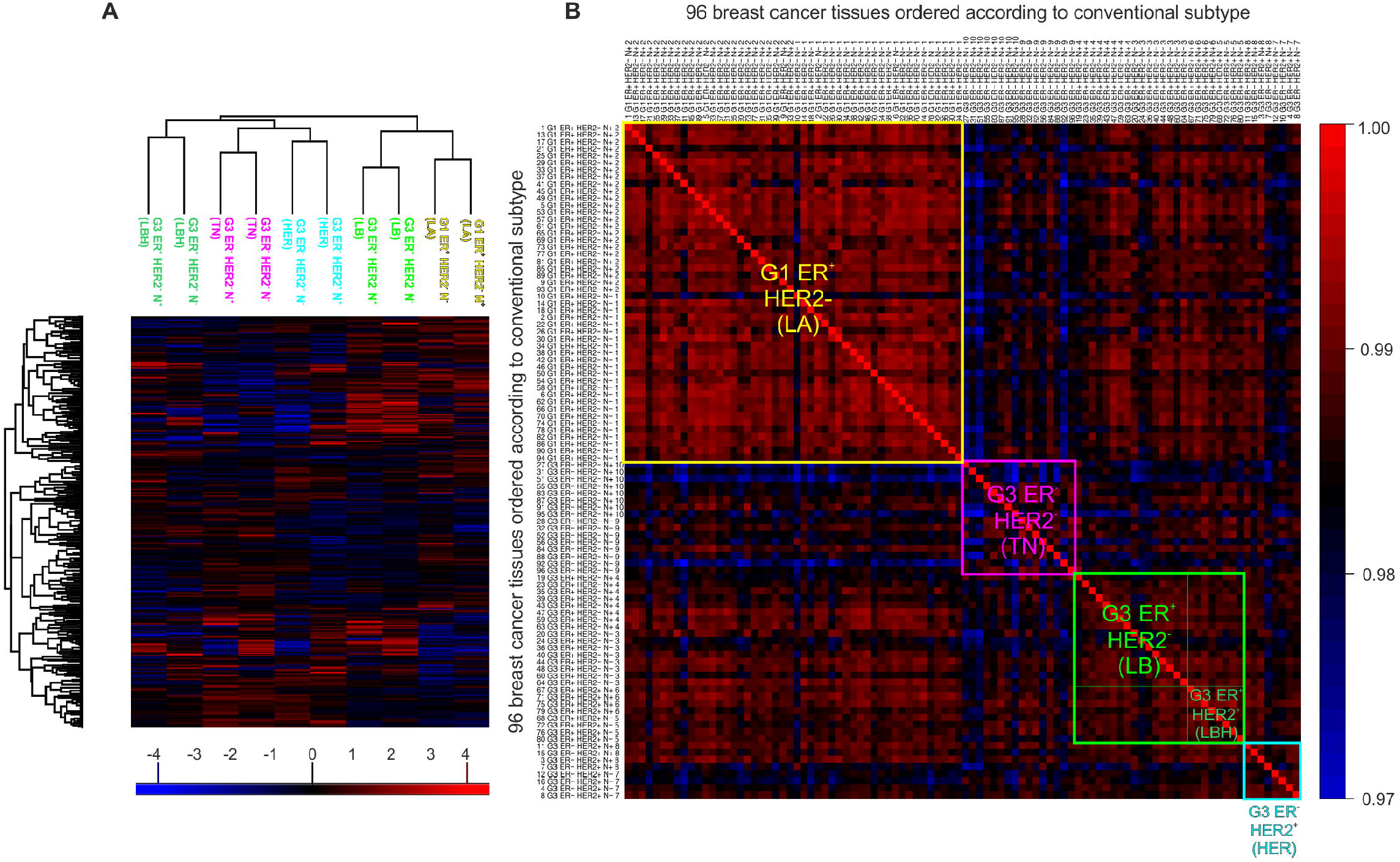
Correlation of breast cancer tissue classification based on conventional subtypes and proteotypes. **(A)** Unsupervised hierarchical clustering of 10 pooled samples from lymph node positive and negative samples of the five conventional breast cancer subtypes. Colours represent log2 protein intensities normalized to median. The sample pools are designated by the conventional subtype nomenclature and colour coded as follows: luminal A (LA, yellow), luminal B (LB, light green), luminal B HER2 positive (LBH, dark green), HER2 enriched (HER, <u>turquoise</u>), triple negative (TN, pink). Lymph node status: negative (N^−^), or positive (N^+^). The figure shows a high similarity of proteotype patterns of pairs of N and N” tissues within each individual subtype. Furthermore, ER positive and HER2 negative subtypes cluster together (see a close clustering of groups designated LA, LB), similarly, ER negative subtypes, HER and TN, cluster together. **(B)** Correlation matrix of 96 individual samples ordered according to their subtypes (see Data file S I for detailed sample number legend). Colours represent correlation of summarized log 2 protein intensities normalized to median, scaled from blue (least correlated) through black to red (most correlated). Correlations of samples within each subtype are visible, most significantly for luminal A and luminal B and for correlation of both these subtypes. Triple negative tumours showthe highest intra-group heterogeneity

Next, we systematically correlated the quantitative proteotypes of the 96 individually measured breast cancer samples and ordered the resulting correlation coefficients according to the classical tumour subtypes (Fig. 1B). Spearman correlation of proteomic profiles across the entire dataset was high (R>0.97). The highest intra-group correlation of proteotypes was within luminal A subtype (R=0.9900). Very high correlation was also observed within the luminal B subtype (both HER2^−^, R=0.9866, and HER2^+^, R=0.9878) and between luminal A and luminal B subtypes (R=0.9865) (Fig. 1B). Furthermore, we found a high correlation of some samples of the HER2 enriched subtype with some (mostly lymph node positive) luminal B HER2^+^ samples, indicating that a higher degree of similarity in HER2^+^ tumours of luminal B and HER2 enriched subtypes were apparent from the proteotype. The group of triple negative tumours exhibited slightly lower inter- (R<0.9852) and particularly intra-group (R=0.9840) correlation, potentially indicating tumour heterogeneity not captured by the conventional tumour classification. In summary, we found that clustering by proteotypes closely recapitulates conventional tumour subtyping, but we also found that some of these subtypes are more heterogeneous (triple negative tumours) than others (Fig. 1B).

### Pathways and proteins associated with key breast cancer characteristics

Having a large high-quality proteomic dataset at hand, we were interested in identifying pathways and proteins that are important for breast cancer biology and progression. We first identified proteins that are differentially expressed in tumours of different ER status, tumour grade, HER2 status or lymph node status (Data file S3). We then used gene set enrichment analysis (GSEA) to find pathways that are enriched among the most differentially abundant proteins in these comparisons (Fig. 2). Among these, there were several pathways known to be associated with the particular phenotype, for example, an enrichment of the NF-κB pathway in ER^+^ tumours, in agreement with its role in proliferation and metastasis of luminal tumours (Azim et al., 2015; Bouchal et al., 2015; Pratt et al., 2009). The list of pathways enriched in high grade tumours was led by the MCM pathway, which includes pro-proliferation proteins of the MCM family regulating cyclin-dependent kinases and DNA replication (Shetty et al., 2005; Wojnar et al., 2010). In HER2^+^ tumours, we found an enrichment of proteins belonging to the VEGF pathway, namely seven up-regulated subunits of Eukaryotic translation initiation factors 2 and 2B, which are known to be regulated by HER2 (Sequeira et al., 2009). In lymph node positive tumours, we found members of the CARM1 and Regulation of the Estrogen Receptor pathway (CARM_ER) to be enriched, potentially indicating an involvement of chromatin remodeling factors in breast cancer progression and metastasis (Wang et al., 2014). All these and further enriched pathways shown in Fig. 2 could be highly relevant for breast cancer biology and warrant further investigation as potential targets of breast cancer therapy.

**Fig. 2.**
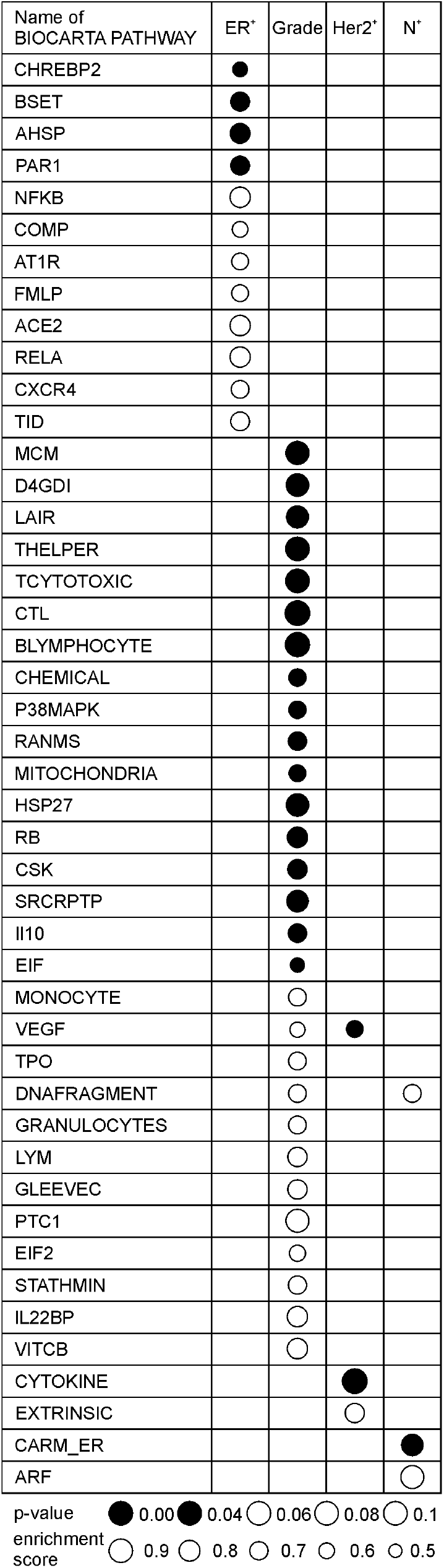
Enrichment of pathways and functional classes among differentially abundant proteins in different tumour phenotypes. Pathway enrichment was performed by gene set enrichment analysis (GSEA) in lists of proteins sorted according to their fold-change in four different comparisons: ER status, HER2 status, tumour grade, and lymph node status. Only pathways enriched in the positive phenotype are shown, i.e. ER positivity, high tumour grade, HER2 positivity and lymph node positivity. Pathways with significance at α=0.1 are displayed and ordered according to p-value.

### Selection of discriminant proteins for improved classification of breast cancer subtypes

To examine the potential of proteotyping for breast cancer classification, we next constructed a decision tree to classify the 96 tumours into the five conventional subtypes based on their proteotypes. We started by selecting the most differentially abundant proteins (log2FC > 1.5, FDR-adj. p-value < 0.05) from the following comparisons: ER^+^ vs. ER^−^ (8 proteins), grade 3 vs. grade 1 (2 proteins), HER2^+^ vs. HER2^−^ (2 proteins), luminal B vs. luminal A (3 proteins), luminal B HER2^+^ vs. luminal A (3 proteins), HER2 enriched vs. luminal A (7 proteins), triple negative vs. luminal A (5 proteins), and HER2 enriched vs. luminal B (2 proteins). This procedure resulted in a list of 22 key proteins (partially overlapping among different comparisons). In a next step, we applied a recursive partitioning algorithm for continuous data in a conditional inference framework (Hothorn et al., 2006). The algorithm automatically selected discriminant proteins from the protein list and provided their quantitative thresholds as well as the structure of the decision tree. The algorithm generated a decision tree with three key nodes (Fig. 3A), representing three key proteins: type II inositol 3,4-bisphosphate 4-phosphatase (INPP4B), cyclin-dependent kinase 1 (CDK1) and receptor tyrosine-protein kinase erbB-2 (ERBB2). Importantly, the differential expression of the selected proteins reflects key clinical parameters defining breast cancer subtypes: ER status (INPP4B, Fig. 3B), tumour grade (CDK1, Fig. 3C) and HER2 status (ERBB2, Fig. 3D). Furthermore, we found that proteotype-based decision tree assigned 84 % of the tumours into their diagnosed conventional subtypes (Fig. 3A).

**Fig. 3.**
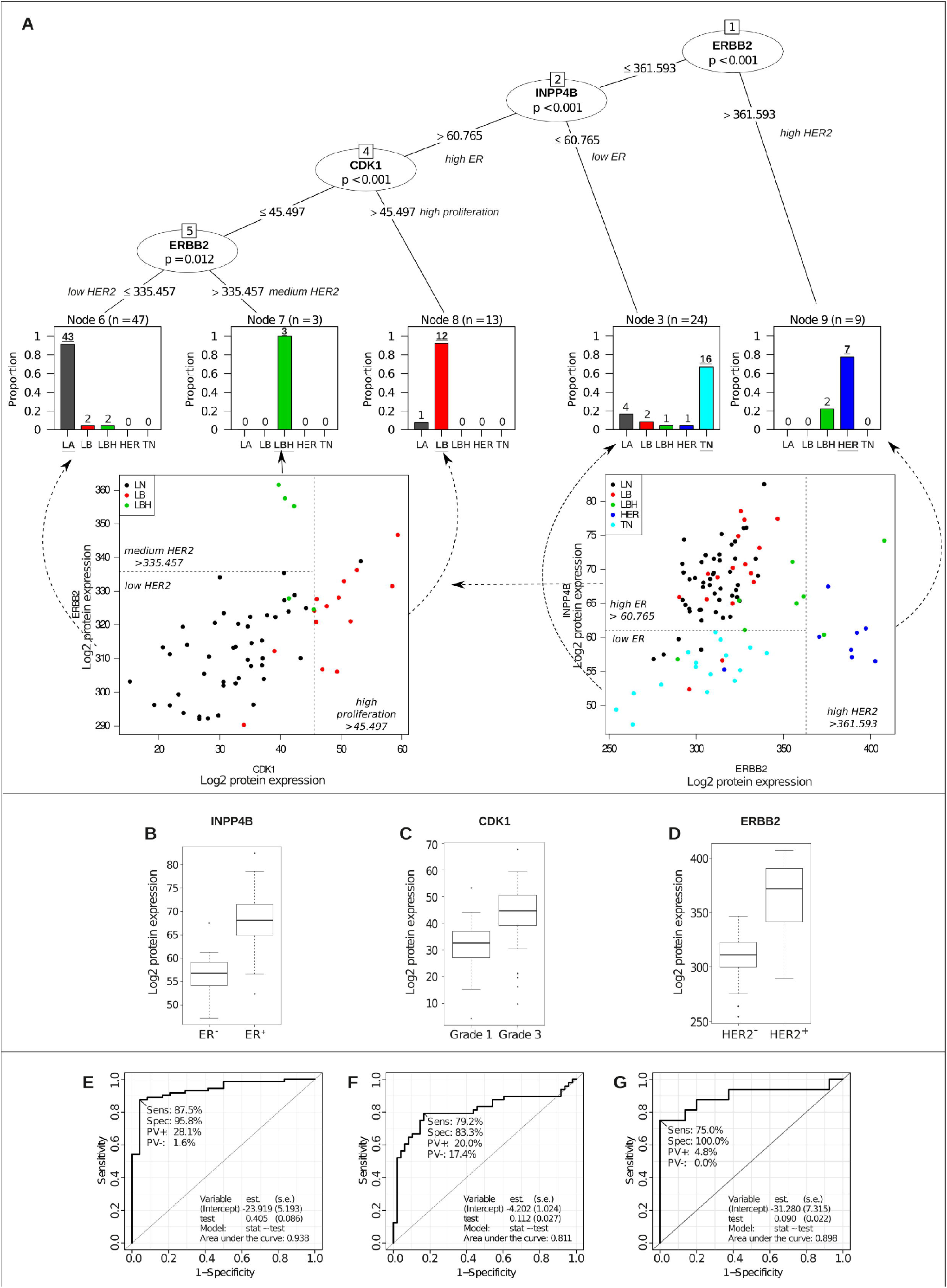
Classification of breast cancer patients based on protein levels in tumour tissue. **(A)** Decision tree classification. The top panel shows the decision tree generated from 22 proteins selected from proteotypes of 96 patients (see Data file S1A-B for details). The bar plots (bottom part of A) show the number of patients, classified by the protein-based decision tree, that coincide with the conventional subtype classification. **(B)-(D)** Protein levels of the classifying proteins are clearly associated with ER status (INPP4B; adj. p=6.73×10^−7^, (**B**)), tumour grade (CDK1; adj. p=1.15×10^−5^; (**C**)) and HER2 status (ERBB2; adj. p=4.55×10^−12^ (**D**)). (E)-(G) ROC curves showing sensitivity and specificity of INPP4B for ER status (**E**), CDK1 for tumour grade **(F)** and ERBB2 for HER2 status **(G)**.

### Validation of the three key proteins selected by the decision tree

We next asked whether the changes in protein levels of the three key proteins from the decision tree, INPP4B, CDK1 and ERBB2, have general discriminative potential and biological validity beyond our 96-patient dataset. Analysis of a published proteomic dataset of 60 human tumour cell lines (http://wzw.tum.de/proteomics/nci60) confirmed high levels of INPP4B protein in ER^+^ breast cancer cell lines (MCF-7 and T47D) while no INPP4B protein was found in ER^−^ breast cancer cell lines (MDA-MB-231, MDA-MB-468, BT549, and HS 578T), supporting the link between INPP4B and ER status. CDK1 and ERBB2 proteins were not covered in this reference dataset. We furthermore compared our protein level data with gene expression data in five published microarray datasets (883 patients, Fig. S3) (Haibe-Kains et al., 2012) and a published RNA sequencing dataset (1078 patients) by The Cancer Genome Atlas (TCGA) (https://portal.gdc.cancer.gov). This analysis confirmed the connection of INPP4B with ER status, CDK1 with tumour grade, and ERBB2 with HER2 status (Fig. 4 and Fig. S4). Furthermore, we found that gene expression of *INPP4B, CDK1, and ERBB2* was statistically significantly connected with patient survival in the same manner as the commonly used reference genes *ESR1* (for ER status) and *MKI67* (for tumour grade/proliferation) (Fig. S5).

**Fig. 4.**
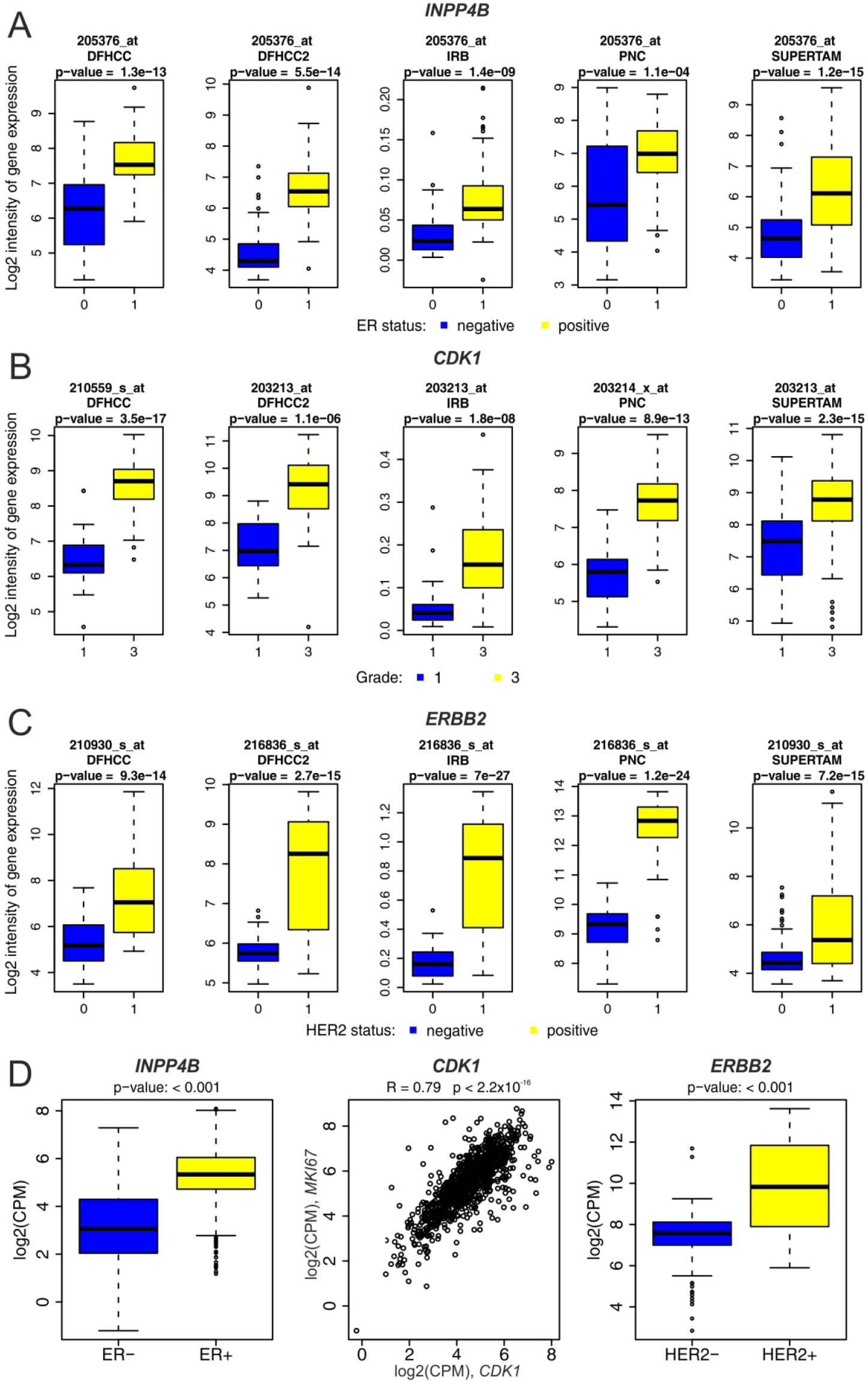
Independent validation of *INPP4B, CDK1* and *ERBB2* association with ER status, tumour grade, and HER2 status. Five independent transcriptomics datasets of 937 patients (DFHCC, DFHCC2, IRB, PNC and SUPERTAM_HGU133PLUS_2 (Haibe-Kains et al., 2012), see Material and Methods and Data file S4D for dataset details) were analysed for gene expression of the three key proteins INPP4B **(A)**, CDK1 **(B)**, and ERBB2 **(C)** (data from the most variable Affymetrix probeset is shown here, see Fig. S4 for data from all probes). For each of the three genes, transcript levels were significantly different (p<0.05) depending on ER status (for *INPP4B*), tumour grade (for *CDK1*), or HER2 status (for *ERBB2*). The Cancer Genome Atlas (TCGA) RNA sequencing dataset of 1078 patients (see Data file S4E for dataset details) **(D)**: transcript levels were significantly different (p<0.05) depending on ER status (for *INPP4B*) or HER2 status (for *ERBB2*); *CDK1* was statistically significantly correlated with proliferation marker *MKI67* (p<0.05).

### Higher level of ERBB2 in ER^−^/HER2^+^ vs. ER^+^/HER2^+^ tumours

An interesting feature of our decision tree is that the algorithm decided between two HER2^+^ subtypes based on ERBB2 protein levels: Whereas lower levels of ERBB2 protein seem to be associated with ER^+^/HER2^+^ grade 3 tumours, higher levels were found in ER^−^/HER2^+^ grade 3 tumours (Fig. 3A). To test whether this observation is of general validity, we manually validated the SWATH-MS-based protein quantification and performed independent analyses at both protein and transcript level. Transcript-level analysis of the same 96 tumour samples described in this study (Bouchal et al., 2015), transcript-level analysis in four additional datasets of a total of 116 tumour samples (Haibe-Kains et al., 2012), as well as immunohistochemistry (IHC) in an independent tumour collection of 78 patients (described in Material and Methods) all confirmed a statistically significantly increased level of ERBB2 in ER^−^/HER2^+^ vs. ER^+^/HER2^+^ tumours (Fig. 5). This observation supports the notion that proteotypes potentially reveal finer graded classification than provided by conventional subtyping.

**Fig. 5.**
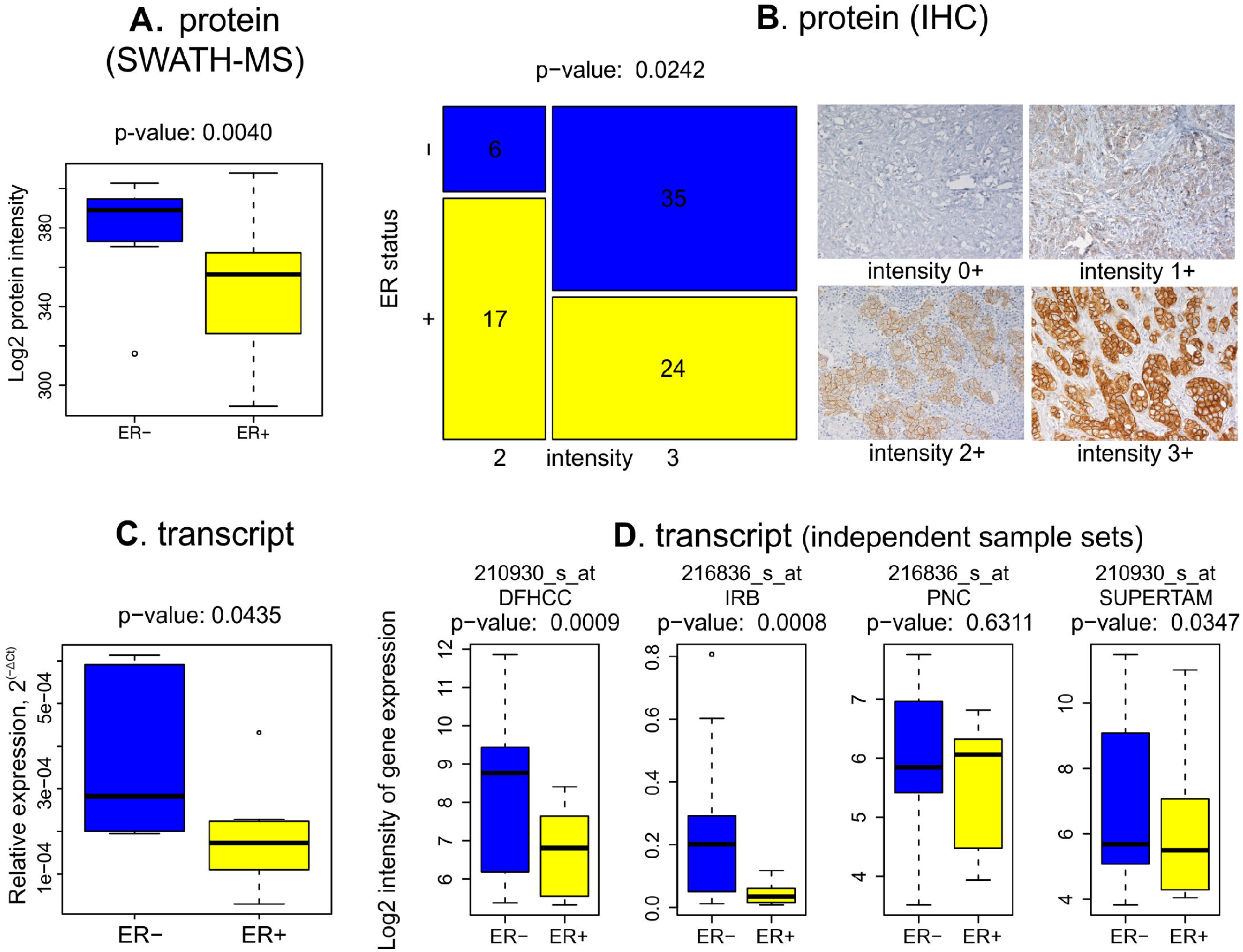
Expression of ERBB2 protein and transcript in ER^−^/HER2^+^ *vs*. ER^+^/HER2^+^ breast cancer tissues. **(A)** Intensity of ERBB2 protein in SWATH-MS proteomics data (16 patients of grade 3, Data file SIB). **(B)** Immunohistochemistry for ERBB2 in an independent set of patients (78 patients of grade 2+3, Data file SIC). **(C)** Transcript-level analysis for ERBB2 in the same patients shown in A (16 patients of grade 3, (Bouchal et al., 2015)). **(D)** Transcript-level analysis in four independent sets of grade 3 patients (DFHCC (27 patients), IRB (23 patients),PNC(24 patients)and SUPERTAM_HGU133PLUS_2(42 patients, DatafileS4D)).

### Analysis of global correlation between proteins and transcripts

To see how our protein-level data correlates with transcript-level data globally, we compared our comprehensive SWATH-MS dataset against the five microarray datasets of 883 patients mentioned above (Haibe-Kains et al., 2012) (see Data file S4 for details). We performed 475,755 individual comparisons of overlaps of differentially abundant proteins (FDR-adj. p-value < 0.05) versus their cognate transcripts (with the same trend) for 2,782 matching transcript-protein pairs between patient groups with different subtype, ER status, HER2 status, tumour grade, and lymph node status (Data file S4). Overall, 6 % of protein-level observations and 7-15 % of transcript-level observations (depending on the set of patients) exhibited statistically significant changes (Data file S4B). Of these, 13-28 % of differentially abundant proteins also showed a statistically significant change with the same direction on the transcript level. From the reverse perspective, 9-18 % of significantly regulated transcripts showed a significant change with the same trend also on protein level. The global correlation coefficients for fold-changes between transcripts and proteins ranged from R=0.17 to R=0.29, depending on the dataset (Fig. 6A). In contrast, the correlation for the three key proteins from the decision tree was very high, with correlation coefficients from R=0.67 to R=0.81 (Fig. 6B). A decision tree constructed from the five independent transcriptomics datasets using expression data for 1036 genes resulted in a tree with three nodes and similar structure (Fig. S3). Taken together, although high correlation of protein and transcript levels was observed for the key proteins INPP4B, CDK1, and ERBB2, correlation and overlap of differentially expressed proteins and transcripts on a global scale was rather low, indicating the importance of protein-level measurements to study breast cancer biology.

**Fig. 6.**
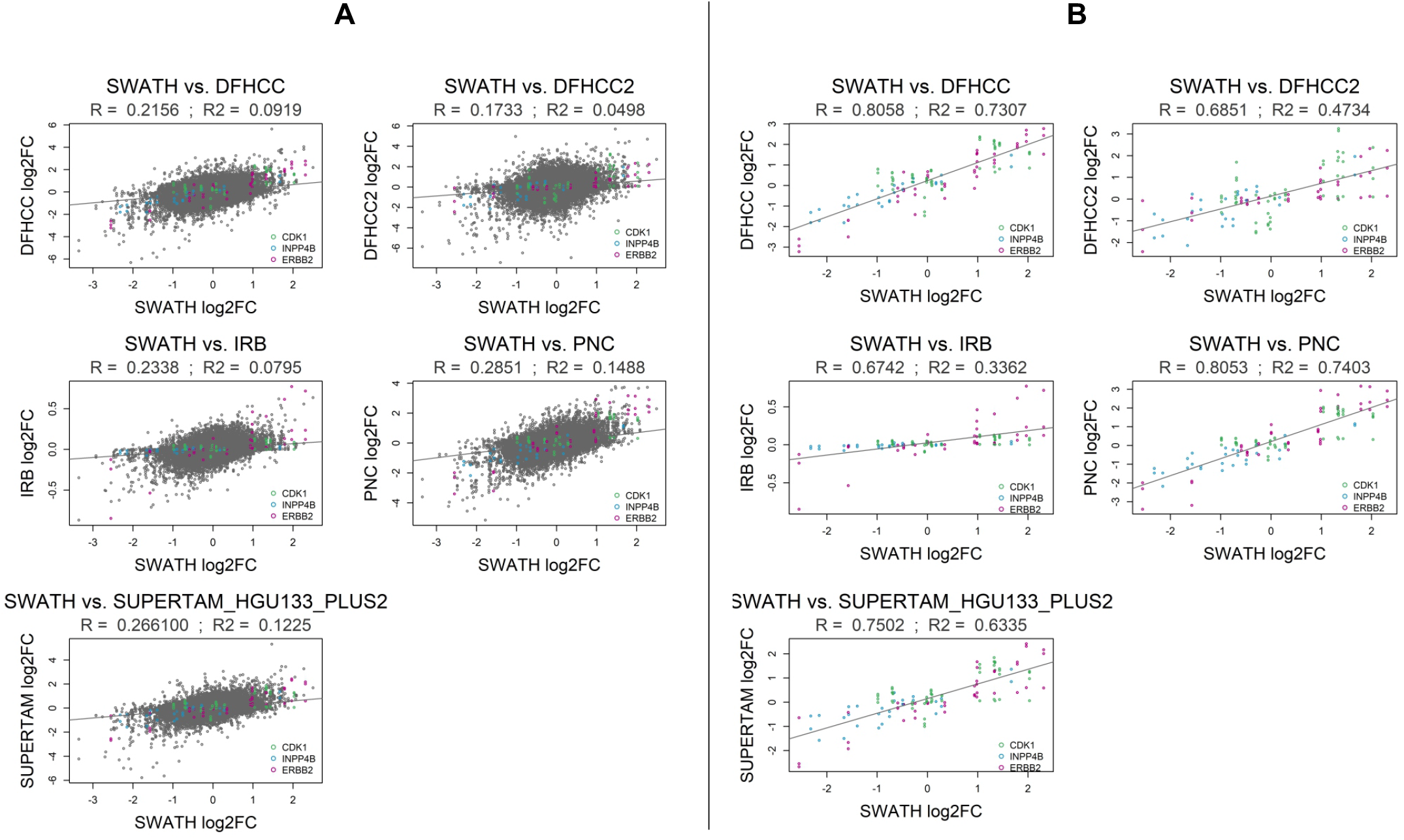
Correlation between protein and transcript levels. Plots show Spearman’s correlation of log2 fold changes (log2FC) between our SWATH-MS protein dataset (96 patients) and five independent transcriptomics datasets DHHCC, DFHCC2, PNC, IRB and SUPERTAM_HGU_133_PLUS2 (Haibe-Kains et al., 2012) (883 patients, see Data file S4 for dataset details) for **(A)** all protein/transcript pairs and **(B)** the three key proteins (vs. transcripts) selected by the decision tree, INPP4B, CDK1, and ERBB2.

## Discussion

### High-throughput proteotyping by SWATH-MS as a next-generation approach for cancer classification

The currently used classification of breast cancer tissues primarily relies on semi-quantitative IHC which is based on manual evaluation of antibody-stained tissue sections by a pathologist. Transcript-level approaches have been used for expression profiling of breast cancer-associated genes and classification, however, as confirmed by our data, gene expression does not generally reflect levels of proteins. Protein-level quantification, although technically more difficult, is hence expected to provide the most relevant information. In this study, we employed for the first time a recently established massively parallel targeted proteomics technique, SWATH-MS, for the classification of human breast cancer tissues. The technique generally requires no more than 1-2 μg of total peptide sample and is capable of analysing tissue samples obtained by needle biopsy (Guo et al., 2015). Moreover, it has good quantitative accuracy with high specificity due to targeted MS/MS data extraction (Gillet et al., 2012), low cost per run, and relatively high sample throughput, enabling the analysis of 10-24 samples per day. The hereby established proteotypes mostly recapitulated the five conventional subtypes, confirming the general applicability of proteotyping for the identification of cancer subtypes. The inconsistencies between the proteotype-based and conventional classification might reflect further breast cancer subtypes (Prat et al., 2015), which could for example arise from additional genetic mutations. This is well illustrated by the *TP53* mutation status in our 96 tumour samples: while 50 % of tumours with more aggressive subtypes (triple negative, HER2 enriched, and luminal B HER2^+^) had mutations in *TP53*, less aggressive luminal B and luminal A subtypes included only 12.5 % and 0.0 % of *TP53*-mutated tumours, respectively (Data file S1B). Proper classification of such additional mutational heterogeneity could help to improve diagnostics and treatment of breast cancer.

### Advantages of SWATH-MS to classify breast cancer tumours

Several studies used proteomics approaches to classify breast cancer tissues (Lam et al., 2014), applying a range of methods, from SELDI-TOF MS (Bouchal et al., 2013; Brozkova et al., 2008) and SILAC-LC-MS/MS (Waldemarson et al., 2016) in breast cancer tumour samples to MS1-based label-free quantification of secreted proteins in a cell line panel (Pavlou et al., 2013). These studies confirm the utility of protein expression profiling for the identification of novel molecular markers to classify breast cancer. We previously analysed the tumour samples of the 96 patients described in the present study using an iTRAQ-2DLC-MS/MS approach in an attempt to identify metastasis-associated proteins in low-grade breast cancer (Bouchal et al., 2015). In that study we quantified 6 % more proteins than in the current study (see also Fig. S1), however, there we were limited by significantly lower sample throughput, only allowing the analysis of pooled and not individual samples in a reasonable time, resulting in inferior statistical power. Compared to the iTRAQ method used earlier by us (Bouchal et al., 2015) and the Clinical Proteomic Tumour Analysis Consortium (Mertins et al., 2016), SWATH-MS has a better quantitative accuracy by avoiding the flattening of peptide ratios due to the use of the same iTRAQ reporter ions for quantification of co-isolated precursors. A recent study using SuperSILAC for the proteomic profiling of 40 breast cancer tissues (Tyanova et al., 2016), identified 10,138 endogenous proteins in total, but only a fraction of this number (2,588 proteins) was quantified across all samples (Fig. S6). The study found a 19-protein signature discriminative for medium- and high-grade breast cancer subtypes, of which we consistently quantified 14 proteins in our SWATH-MS dataset of 96 patients. The abundance ranks of these 19 proteins in the two independent datasets (their 40 patients and our 96 patients) was highly similar (Tab. S1). Compared to the SuperSILAC approach, advantages of SWATH-MS are the lower cost and convenience of the label-free quantification, but most importantly, the consistent quantification of proteins across large sample sets (Fig. S6). One of the gold-standard methods to profile proteins in clinical tissue samples is selected/multiple reaction monitoring (S/MRM). Of 319 breast cancer-associated proteins quantified by S/MRM by Kennedy and colleagues (Kennedy et al., 2014) our SWATH-MS data covers 305 (96 %). Similarly, 9 of 10 proteins associated with breast cancer biology (represented by 16 of 17 peptides) were quantified by S/MRM in the same set of tumours (Prochazkova et al., 2017) as in our current SWATH-MS data set with high level of correlation (Spearman correlation coefficients 0.439 to 0.880 and p-values 1.1*10^−5^ to 2.2*10^−16^, Data file S5). This comparison well validates our SWATH-MS data using an independent method on individual tumour level. A strong correlation between SWATH-MS and S/MRM was demonstrated already in the first SWATH-MS publication (Gillet et al., 2012) and confirmed in other independent studies (Kockmann et al., 2016; Liu et al., 2013; Nakamura et al., 2016; Schmidlin et al., 2016). These studies include our recent comparison of S/MRM, pseudo-SRM/MRM^HR^ and SWATH-MS analytical parameters in selected samples from the same breast cancer tissue collection (Faktor et al., 2017). Based on the above data, it has been well demonstrated both experimentally in our breast tumour sample set and in the literature that SWATH-MS provides data highly correlated with S/MRM. In summary, our SWATH-MS-based strategy provided an advantageous combination of sample throughput, quantitative precision (Vowinckel et al., 2013), and proteome coverage in a large sample set. Applying the latest technical developments (e.g., ion mobility MS, faster Orbitrap-based instruments) may further improve the quantitative depth of SWATH-MS or similar data-independent acquisition-based studies.

### Biological relevance of the key proteins selected by the decision tree

The three key proteins identified by our decision tree are strongly associated with important clinical parameters, namely oestrogen receptor status (INPP4B), tumour grade (CDK1), and HER2 status (ERBB2) (Fig. 3B-3D). The receptor tyrosine protein kinase ERBB2 is the protein product of the *HER-2/NEU* gene and is routinely being used for breast cancer classification into HER2^+^ or HER2^−^ phenotypes. It also is the target of anti-HER2 therapy via US Food and Drug Administration (FDA)-approved humanized monoclonal antibody trastuzumab. The strong association of the ERBB2 protein with HER2 status in our dataset internally validates our proteomics data and design of the study. Of note, higher levels of ERBB2 protein observed here in ER^−^/HER2^+^ vs. ER^+^/HER2^+^ tumours are also consistent with the better response to therapy of ER^−^/HER2^+^ vs. ER^+^/HER2^+^ tumours (Bhargava et al., 2011).

INPP4B is known to dephosphorylate phosphatidylinositol 3,4-bisphosphate in the PI3K pathway, which co-activates cell growth and movement via Akt kinases (Malek et al., 2017). Hence, it serves as a tumour suppressor and our and earlier observations that it is significantly associated with ER^+^ tumours (Fedele et al., 2010) suggest that it should be explored as a candidate therapeutic target for ER^+^ breast cancer. The second of our key proteins, mitotic kinase CDK1, is known to accelerate critical processes required for mitosis (Enserink and Kolodner, 2010) and correlates with tumour grade (Chae et al., 2011). Moreover, inhibitors of the family members CDK4/6 have been FDA-approved for the treatment of metastatic breast cancer in a first-line setting (Bilgin et al., 2017). In conclusion, although this is a pilot discovery study and follow-ups with larger patient cohorts are required to further train and validate our classifier, our findings suggest that both INPP4B and CDK1 are promising alternative targets for anti-cancer therapy, as they exhibit similar level of association with ER status and tumour grade as ERBB2 with HER2 status, which is already successfully targeted to treat HER2^+^ breast cancer patients. We would like to note that subsequent validation studies with S/MRM can now be set up easily as the information needed to the required acquisition methods can be obtained directly from the SWATH assay library (see also Data file S6). A small panel of validated protein biomarkers could be subsequently implemented as part of an IHC panel or assessed with other techniques used in the clinic.

### Molecular features available in proteotype and not in conventional subtype

Although there was a high concordance (84 %) between classification based on proteotypes and conventional subtypes, some samples with identical conventional subtype showed distinct proteotypes. We find such proteotype heterogeneity for example in triple negative tumours, a genetically heterogeneous group that can indeed be sub-divided further. For example, Lehmann et al. suggested six subtypes based on gene expression profiling: basal-like 1, basal-like 2, immunomodulatory, mesenchymal, mesenchymal stem-like, and luminal androgen receptor subtype (Lehmann et al., 2011); others suggested a similar division (Palma et al., 2015). We also found that some HER2-enriched tumours were more similar to luminal B HER2^+^ tumours than reflected in current subtyping; this is evident also in data from Brozkova and colleagues (Brozkova et al., 2008). All these data indicate that proteotypes have the potential of enabling finer stratification of a patient population than conventional subtyping. Current clinical practice shows that treatment based on conventional subtypes is far from optimal with respect to patient response, and proteotyping can potentially provide a more accurate picture of the actual molecular state of a cancerous tissue and could thereby enable more precise or even personalized treatment.

### Global correlation and overlaps of protein and transcript level expression

Abundance of our three key proteins INPP4B, CDK1 and ERBB2 across tumours of different ER status, tumour grade, and HER2 status correlated well with their respective transcript levels. However, when looking at all differentially expressed proteins and transcripts in our dataset, the overlap and correlation of fold-changes was modest. While it has been shown that protein levels are chiefly determined by transcript levels, particularly in steady state (Schwanhausser et al., 2011), and that fold-changes of transcript and protein levels between different human cell lines can show correlations as high as R=0.63 (Lundberg et al., 2010), our comparisons of transcript and protein data suggests that this correlation is relatively low in human breast cancer tissues (R=0.29). In general, the limited correlation between protein and transcript levels provides a substantial reason to focus on the analysis of proteins instead of transcripts as these represent the true molecular effectors in cells.

### Conclusions

This study explored and confirmed the suitability of SWATH-MS for proteotyping of human tumour samples at relatively high throughput. While larger patient cohorts are needed for validation, our results indicate that proteotype-based classification resolves more breast cancer subtypes than apparent from conventional subtyping and potentially improves current classification which in turn may result in more adequate treatment and better clinical outcomes. The breast cancer SWATH assay library and the high-quality proteomics dataset of 96 breast cancer tumours will provide a valuable resource for future protein marker studies.

## Acknowledgments

We thank the women who provided their tissue for this research and all of the clinically related staff involved in their treatment. We are grateful to Dr. Hannes Röst and Dr. George Rosenberger for their help with SWATH-MS data analysis, to Dr. Ben Collins for the script to generate Fig. S8, to Dr. Josef Planeta for nano-LC column preparation and to Dr. Philip J. Coates for critical reading of the manuscript. This work was supported by Czech Science Foundation (project No. 17-05957S), J.F. was supported by MEYS - NPS I - LO1413, R.N. by MH CZ - DRO (MMCI, 00209805), E.B. by the CETOCOEN PLUS project and the RECETOX Research Infrastructure (LM2015051). R.A. was supported by the Swiss National Science Foundation (SNSF; 31003A_130530) and European Research Council grant 20140AdG 670821.

## Author contributions

P.B. supervised SWATH-MS analyses at MMCI, analysed SWATH-MS data, coordinated the study and wrote the paper; O.T.S. supervised SWATH-MS data analysis and wrote the paper; J.F. performed SWATH-MS analyses at MMCI; L.C. performed data analysis-clustering and comparison of gene expression at proteome and transcriptome level as well as GSEA pathway analyses; H.I. performed validation of selected gene products in independent data sets; K.Z. performed KEGG pathway analyses; V.P. performed SWATH-MS data analysis in Skyline software; R.H. analyzed p53 status of tumours; Y.L. significantly contributed to SWATH-MS measurements and to manuscript preparation; H.A.E. contributed to SWATH-MS measurements and to manuscript preparation; E.B. constructed the decision tree and co-supervised all data analyses; R.N. designed and selected the set of tissues and contributed to data interpretation; R.A. approved the joint study, provided computational and instrument capacity, wrote and approved the manuscript.

## Materials and Methods

### Study design

The objective of the study was to compare classification of breast cancer tissues based on proteotypes obtained using a novel next generation proteomics approach, SWATH-MS, with clinically used subtypes classified by immunohistological markers and grade. To avoid lymph node status as confounding factor in tumour classification into subtypes, we decided for the same representation of lymph node positive and lymph node negative tumours in the sample set. A secondary aim was to compare SWATH-MS data with previous measurement using discovery proteomics via data dependent analysis (DDA) in the same sample set. To this end we designed a retrospective pilot discovery study on a cohort of well characterized breast tumour samples available from Masaryk Memorial Cancer Institute (MMCI) (Bouchal et al., 2015). The samples were analysed by SWATH-MS and the findings confirmed both by immunochemical validation and by meta-analysis of corresponding mRNA levels in independent publicly available sets of patients.

### Tissue procurement and patient characteristics

Informed patient consent forms and the use of collected tissues for targeted proteomics analysis were approved by the Ethics committee of the Masaryk Memorial Cancer Institute (MMCI). Breast cancer tissue samples were frozen in liquid nitrogen within 20 minutes of surgical removal and stored at −180°C in the tissue bank at MMCI. A set of 96 preoperatively untreated breast carcinomas of 11-20 mm maximum diameter (pT1c) was selected. The set consisted of 48 ER^+^, PR^+^, HER2^−^, grade 1 tumours (luminal A (LA) subtype); 16 ER^+^, PR^+/-^, HER2^−^, grade 3 tumours (luminal B (LB) subtype); 8 ER^+^, PR^+/-^, HER2^+^, grade 3 tumours (luminal B-like HER2 positive (LBH) subtype); and 16 ER^−^, PR^−^, HER2^−^, grade 3 tumours (triple negative (TN) subtype). Half of the tumours in each group was lymph node positive and half was lymph node negative at the time of diagnosis. Full details are available in Data file S1A-B. The cases were reviewed by involved pathologist (Rudolf Nenutil) before entering the study. The tumours were classified and reviewed using FFPE blocks, taken in parallel with the native deeply frozen samples used for proteomics. The samples with very low cellularity of invasive tumour component (e.g. below 20 %), and/or dominant in-situ component and/or apparent clonal morphological heterogeneity were not used. As the dataset attempted to represent the main phenotypes, the cases were of variable malignancy and different cellularity. On average, the low grade tumours are inherently of lower cellularity compared to high grade ones. Based on the results, additional independent set of 78 grade 2 and 3 breast tumours was used for IHC validation of ERBB2 protein levels in HER2^+^, ER^+^ (N=41) vs. HER2^+^, ER^−^ (N=37) tumours (Data file S1C). Sample sets used for meta-analysis of mRNA levels are described in Statistical analysis section.

### TP53 sequencing

Total cellular RNA was extracted using TRI Reagent (MRC). *TP53* mRNA from tumour tissue was amplified using the SuperScript™ III One Step RT-PCR System with Platinum^®^ Taq High Fidelity (Invitrogen), sense primer: 5’ TCCCCTCCCATGTGCTCAAGACTG 3’and antisense primer: 5’ GGAGCCCCGGGACAAAGCAAATGG 3’. PCR products were purified by MinElute™ PCR Purification Kit (Qiagen) and sequenced using the ABI PRISM BigDye^®^ Terminator v 3.1 Cycle Sequencing Kit on an ABI 3130 genetic analyser (Applied Biosystems).

### Tissue quality control via RNA integrity measurement

After homogenization in a MM301 mechanical homogenizer (Retsch, Haan, Germany) using a metal ball for 2×2 min at 25 s^−1^ in 600 μl of RLT buffer (Qiagen, Germany) with 1% β-mercaptoethanol, total RNA was isolated using RNeasy Mini Kit (Qiagen, Germany) following the manufacturer’s protocol. RNA was eluted with 30 μl of RNase-free water, quantified at 260 nm using NanoDrop ND-1000 (Thermo Fisher Scientific, USA) and quality checked by measurement of RNA integrity number (RIN) on Agilent 2100 Bioanalyzer (Agilent Technologies, USA). Samples which did not pass the criterion of RNA quality (RIN > 7) were excluded and replaced by other tissues with the same clinicopathological characteristics for the SWATH-MS experiments.

### Proteomics sample preparation

Frozen breast cancer tissue (approx. 20 mm^3^) was homogenized in 150 μl lysis buffer (6 M guanidine hydrochloride; 0.1 M Na-phosphate buffer, pH 6.6; 1% Triton X-100) in a MM301 mechanic homogenizer (Retsch, Germany) using a metal ball for 2 × 2 min at 20 s^−1^, needle-sonicated (Bandelin 2200 Ultrasonic homogenizer, Bandelin, Germany; 30 × 0.1 s pulses at 50 W) and kept on ice for 1 h. After 14,000 x g centrifugation at 4°C for 20 min, protein concentration was measured in the supernatant using RC-DC assay (Bio-Rad, USA). An aliquot of the lysate containing 60 μg total protein mass was digested using a filter aided sample preparation protocol (Wisniewski et al., 2011) with modifications. Briefly, aliquots of the lysate were mixed with 200 μl 8 M urea in 0.5 M triethylammonium bicarbonate (TEAB) pH 8.5 on Vivacon 500 filter device, cut-off 10K (Sartorius Stedim Biotech GmbH, Germany). The device was centrifuged at 14,000 × g at 20°C for 20 min (all of the following centrifugation steps were performed applying the same conditions). Subsequently, 100 μl 5 mM tris(2-carboxyethyl)phosphine in 8 M urea, 0.5 M TEAB, pH 8.5 was added to the filter, proteins were reduced at 37°C for 60 min at 600 rpm and centrifuged. Next, 100 μl 10 mM S-methyl methanethiosulfonate in 8 M urea and 0.5 M TEAB, pH 8.5 were added to the filter, cysteine groups of peptides were alkylated at 20°C for 10 min and centrifuged. The resulting concentrate was diluted with 100 μl 8 M urea in 0.5 M TEAB, pH 8.5 and concentrated again. This step was repeated twice. The concentrate was subjected to proteolytic digestion by adding 100 μl 0.5 M TEAB, pH 8.5 containing trypsin (TPCK treated, SCIEX, USA) reconstituted in water (trypsin to protein weight ratio 1:30) and by incubating at 37°C for 16 h. The digests were collected by centrifugation into clean tubes, dried in a vacuum concentrator and C18 desalted as previously described (Bouchal et al., 2009) using 0.1% trifluoracetic acid as an ion pairing reagent. Eleven retention time anchor peptides (commercial iRT peptide solution, Biognosys, Zurich, Switzerland) (Escher et al., 2012) were added into each sample at a ratio of 1:40 v/v. For SWATH-MS analysis, equal amounts of samples (estimated to be 1.33 μg protein) were injected in single technical replicates.

### LC-MS analyses for spectral library generation

As an input for generating the SWATH-MS assay library, the following samples were prepared: (i) 10 pooled samples (each pooled from 4-8 patients) of 5 the breast cancer subtypes mentioned above. Each subtype group involved two pools of tumours (lymph node positive and lymph node negative cases separately); (ii) pool of aliquots of all samples in the sample set (400 μg in total) fractionated using HILIC chromatography as follows: HILIC Kinetex column (Phenomenex, USA, 2.6 μm, 150 x 2.1 mm, 100 A) was run in an Agilent Infinity 1260 LC system (Agilent, USA). Mobile phase (A) was composed of 100% acetonitrile (Merck, Germany), mobile phase (B) of water (MilliQ, Millipore) and mobile phase (C) of 50 mM ammonium formate (pH 3.2). 20 μL mobile phase (B) were added to the sample which was then sonicated in an ultrasonic bath for 2 min. Then, 20 μL mobile phase (A) and 5 μL mobile phase (C) were added. After a further 2 min of sonication, the sample was centrifuged at 16,000 x g at 20 °C for 20 min. The sample injection volume was 40 μL and the separation method was set as follows: 5 min isocratic 0% B, 7 min gradient to 20 % B, 23 min gradient to 34 % B, 5 min gradient to 50 % B, 5 min isocratic 50 % B, 0.5 min gradient to 0 % B and for 4.5 min isocratic 0 % B; mobile phase C was kept at 10 % all the time. The flow rate was 0.2 mL/min, column temperature was set to 30 °C and the UV signal was monitored at 280 nm. Fractions were collected every 1 min, some neighbouring fractions with lower signal intensity were subsequently pooled to generate a final set of 20 fractions with similar peptide content. These were vacuum-dried and stored at −80°C.

MS/MS datasets for spectral library generation were acquired on a TripleTOF 5600^+^ mass spectrometer (SCIEX, Canada) interfaced to an Eksigent Ekspert nanoLC 400 system (SCIEX, Canada). Prior to separation, the peptides were concentrated on a C18 PepMap100 pre-column (Thermo Fisher Scientific, USA; particle size 5 μm, 100 Å, 300 μm x 5 mm). After 10 min washing with a solvent consisting of 2 % acetonitrile and 0.0 5% (v/v) trifluoroacetic acid, the peptides were eluted from a capillary column (75 μm × 250 mm, X-Bridge BEH C18 130 Å, particle size 2.5 μm, Waters, USA, prepared as described in (Planeta et al., 2003)) using 2 % mobile phase B for 10 min (mobile phase A was composed of 0.1 % (v/v) formic acid in water, mobile phase B of 0.1 % (v/v) formic acid in acetonitrile) followed by gradient elution from 2 % to 40 % mobile phase B in the next 120 minutes at a flow rate of 300 nl/min. Output of the separation column was directly coupled to nano-electrospray source. MS1 spectra were collected in the range of 400–1250 m/z for 250 ms. The 20 most intense precursors with charge states of 2 to 5 that exceeded 50 counts per second were selected for fragmentation, rolling collision energy was used for fragmentation and MS2 spectra were collected in the range of 200-1600 m/z for 100 ms. The precursor ions were dynamically excluded from reselection for 12 s. All MS/MS data files in wiff and mzXML format are available at http://www.peptideatlas.org/PASS/PASS00857 (reviewer password: BrCa).

### LC-MS analysis in SWATH-MS mode

SWATH-MS datasets of the individual patients were acquired on a TripleTOF 5600+ mass spectrometer (SCIEX, Canada); the same chromatographic system, settings and gradient conditions as described above for spectral library generation were used. Using an isolation width of 9.7 m/z (containing 1 m/z for the window overlap), a set of 69 overlapping SWATH windows was constructed covering the precursor mass range of 400-1000 m/z. The effective isolation windows can be considered as 400.5-408.2 (first narrower window), 408.2-416.9, 416.9-425.6 etc. SWATH MS2 spectra were collected from 360 to 1460 m/z. The collision energy was optimized for each window according to the calculation for a charge 2^+^ ion centered upon the window with a spread of 15 eV. An accumulation time (dwell time) of 50 ms was used for all fragment ion scans in high-sensitivity mode, and for each SWATH cycle a survey scan was also acquired for 50 ms, resulting in a duty cycle of 3.5 s and a typical LC peak width of ~30 s.

Compared to the above conditions, for the analysis of pooled samples (see previous paragraph for pooling scheme) the parameters were changes as follows: (i) chromatographic separation of peptides was performed on 20-cm emitter (75 μm inner diameter, #PF360-75-10-N-5, New Objective) packed in-house with C18 resin (Magic C18 AQ 3 μm diameter, 200 Å pore size, Michrom BioResources); (ii) a linear gradient from 2-30 % solvent B (98 % ACN/0.1 % FA) was run over 120 min at a flow rate of 300 nl/min; (iii) because of the increased sample complexity due to the pooling strategy, a set of 64 SWATH windows (containing 1 m/z for the window overlap) with variable width optimized for human samples was used to cover the precursor mass range of 400-1200 m/z (Collins et al., 2017). All SWATH-MS raw data are available at http://www.peptideatlas.org/PASS/PASS00864 (reviewer password: BrCa).

### ERBB2 immunohistochemistry

After removal of paraffin wax with xylene and rehydration, endogenous peroxidase activity was blocked with 3 % hydrogen peroxide in phosphate buffered saline (PBS) pH 7.5, for 15 minutes. No antigen retrieval was performed. After three PBS washes, nonspecific binding activity was blocked with 5 % non-fat dried milk in PBS for 15 minutes. The cocktail of anti HER-2 primary antibodies was diluted in antibody diluent (DakoCytomation, Denmark A/S) to 1:500 for Novocastra NCL-c-erbB-2-316, and 1:1,000 for Novocastra NCL-L-CBE-356 (both Leica Biosystems) and applied overnight at 4°C. Reactive sites were identified with biotinylated antimouse and anti-rabbit secondary antibodies and peroxidase ABC reagents (Vector-Elite, Vector Laboratories, Burlingame, CA, USA) according to the manufacturer’s instructions and peroxidase activity was visualized with DAB+ reagents (DakoCytomation). Sections were washed in distilled water and counterstained with Gills haematoxylin, dehydrated, cleared and mounted. Membrane staining of tumour cells was evaluated as 0, 1+, 2+, 3+ according to the HercepTest™ Interpretation Manual (DakoCytomation).

### Data processing

#### a. SWATH-MS assay library generation

Raw data files (wiff) were centroided and converted into mzML format using the SCIEX converter (beta release 111102) and subsequently converted into mzXML using openMS (version 1.9.0, Feb 10 2012, Revision 9534). The converted data files were searched using the search engines X!Tandem (k-score, version 2011.12.01.1) and Comet (version 2013.02, revision 2) against all human proteins annotated in UniProt/SwissProt (2014_04) and the sequences of 11 iRT peptides (iRT-kit, Biognosys). The searched database also contained a decoy protein sequence (reversed protein sequence) for each database protein. Only fully tryptic peptides with up to two missed cleavages were allowed for the database search. The tolerated mass errors were 15 ppm on MS1 level and 0.1 Da on MS2 level. Methylthiolation of cysteines was defined as a fixed modification and methionine oxidation as a variable modification. The search results were processed with PeptideProphet (Keller et al., 2002) and iProphet (Shteynberg et al., 2011) as part of the TPP 4.6.0 (Deutsch et al., 2010). The SWATH-MS assay library was constructed from the iProphet results with an iProphet cut-off of 0.8360 which corresponds to 1% FDR on peptide level. The raw and consensus spectral libraries were built with SpectraST (version 4.0) (Lam et al., 2007; Lam et al., 2008) using the -cICID_QTOF option for high resolution and high mass accuracy. Retention times were normalized and converted to iRT space using spectrast2spectrast_iRT.py (imsproteomicstools R356). The 6 most intense y and b fragment ions of charge state 1, 2 and 3 were extracted from the consensus spectral library using spectrast2tsv.py (imsbproteomicstools). Neutral losses −17 (NH_3_), −18 (H_2_O) and −64 (CH_4_SO, typical for oxidized methionine) were also included if they were among the 6 most intense fragment ions. Fragment ions falling into the SWATH window of the precursor were excluded as the resulting signals are often highly interfered. The library was converted into TraML format using the OpenMS tool ConvertTSVToTraML (version 1.10.0). Decoy transition groups were generated based on shuffled sequences (decoys similar to targets were excluded) by the OpenMS tool OpenSwathDecoyGenerator (version 1.10.0) and appended to the final SWATH library in TraML format. All intermediary and final files of the library building workflow are available at http://www.peptideatlas.org/PASS/PASS00857 (reviewer password: BrCa).

#### b. SWATH-MS data processing in OpenSWATH

The SWATH-MS data was analysed using OpenSWATH (Rost et al., 2014) with the following parameters: Chromatograms were extracted with 0.05 Th around the expected mass of the fragment ions and with an extraction window of +/-5 min around the expected retention time (see Data file S3C for justification). The best models to separate true from false positives (per run) were determined by pyProphet with 10 cross-validations. The runs were subsequently aligned with a target FDR of 0.01 for aligned features (Rost et al., 2016). Background signals were extracted for features that could not be confidently identified (Rost et al., 2016). To reduce the size of the output data and remove low-quality features, two filtering steps were introduced: (i) keep only the 10 most intense peptide features per protein and (ii) of these, keep only features that were identified with an FDR<0.01 in at least four samples over all runs, corresponding to the smallest tumour group in the dataset defined by a combination of subtype and lymph node status. The OpenSWATH output files are available at http://www.peptideatlas.org/PASS/PASS00864 (reviewer password: BrCa).

### Statistical analysis

All statistical tests were two-tailed and the results were considered statistically significant at alpha=0.05 or FDR=0.05, if not stated otherwise. Definition of error bars in all figures: Boxes are extended from the 25th to the 75th percentile, with a line at the median. The whiskers extend to the most extreme data point which is no more than 1.5 times the interquartile range (IQR) from the box. The individual points represent outliers or extreme values.

#### a. Relative quantification with MSstats and differential protein expression analysis between subtypes and related clinical-pathological variables

We used the R (version 3.0.3) package MSstats 2.1.3 (Choi et al., 2014) for relative quantification of protein levels among the five different breast cancer conventional subtypes and related clinical-pathological variables (ER, grade, HER2, lymph node status). Before MSstats and correlation analysis, the OpenSWATH output was further reduced to contain up to five peptide features per protein and the intensities were log2 transformed and median-equalized. The differences in protein expression between conventional subtypes and related clinical-pathological variables were compared pairwise using mixed effect models as implemented in the `groupComparison` function of MSstats, with expanded scope of biological and restricted scope of technical replication. Resulting p-values were corrected for multiple hypotheses testing by the Benjamini-Hochberg method.

#### b. KEGG pathway analysis

The list of 4,443 proteins in the SWATH-MS library of assays (Data file S2A) and the list of SWATH-MS 2,842 quantified proteins (Data file S3) were inserted in Kyoto Encyclopedia of Genes and Genomes (KEGG) Mapper (www.kegg.jp/kegg/tool/map_pathway2.html), searched against hsa (Homo sapiens) database the subset of proteins related to Pathways in cancer (hsa05200) was displayed.

#### c. Gene set enrichment analysis

Gene set enrichment analysis (GSEA) in GSEA Java desktop application (http://software.broadinstitute.org/gsea/downloads.jsp) was conducted using the pre-ranked list (according to protein fold changes between ER^+^/ER^−^, tumour grade 3/grade 1, HER2^+^/HER2^−^, lymph node positive/negative patient groups) of 2,842 proteins quantified by SWATH-MS to find pathways enriched in ER^+^, high grade, HER2^+^, and lymph node positive phenotypes separately, with a priori defined pathways from BioCarta (https://cgap.nci.nih.gov/Pathways/BioCarta_Pathways). We used default settings, except that we decreased the minimal size of a gene set to 1 and we did not use any normaliation method to normalize the enrichment scores across analysed gene sets.

##### d. Correlation analysis of breast cancer tissue proteomes

For the correlation analysis of the pooled samples, label-free quantification was conducted using the R package aLFQ (1.3.2) (Rosenberger et al., 2014). The method `ProteinInference` with default parameters (summing the three most intense transitions per peptide and averaging the two most intense peptides per protein) but without consensus feature selection was used to compute a protein intensity for all 1,832 proteins for which at least one peptide has been quantified by OpenSWATH (only including proteotypic peptides). Hierarchical clustering with Spearman’s correlation-based distance matrix and average linkage algorithm was performed in Perseus 1.5.1.6 software (www.maxquant.org) on log2 transformed, Z-score normalized (on both samples and proteins according to median) protein abundance values, including only proteins quantified in all pools. For correlation analysis of individual samples, we selected all 2,842 proteins for which proteotypic peptides were quantified by OpenSWATH and performed Spearman’s correlation among samples based on log2 protein intensities.

##### e. Construction of the decision tree

We used a conditional reference tree algorithm for automated selection of the most discriminative variables (proteins) between conventional subtypes and for the generation of a decision tree, using ctree() function of R package party (Hothorn et al., 2006) with the control parameters set to default, except for the minimum sum of weights in a node in order to be considered for splitting (minsplit = 50) and the minimum sum of weights in a terminal node (minbucket=3). The analysis was based on the set of 22 proteins with significantly different abundances between different conventional subtypes and related clinical-pathological characteristics, as determined by MSstats and presented in the Results part. The selected proteins were further validated in gene expression datasets and through immunohistochemistry. The decision tree was constructed also on gene expression data (see section *g* below).

##### f. Analysis of ERBB2 gene expression in the same sample set

The data were extracted from our previously published dataset (Bouchal et al., 2015).

##### g. Analysis of gene expression in independent microarray and RNA-Seq sets of samples

Publicly available gene expression datasets DFHCC, DFHCC2, IRB, PNC and SUPERTAM_HGU133PLUS_2 (all platform Affymetrix Human Genome U133A, 937 samples in total) were downloaded in the log2 normalized form (Haibe-Kains et al., 2012) and used in order to confirm at the transcriptome level the hypotheses derived from our analysis of proteomic SWATH-MS data. For this purpose, subsets of 883 samples with available information on gene expression, ER status, tumour grade, HER2 status or lymph node status were used, based on the type of comparison (see Fig. S2).

First, we performed analysis of differential expression between conventional subtypes (pairwise) and between the categories of clinical-pathological variables using moderated t-statistics (method (Choi et al., 2014) implemented in the R package Limma of R 3.0.2), on the set of 6,895 probesets representing 2,782 genes with corresponding products (proteins) measured also by SWATH-MS in our experiment. This means that out of 2,842 proteins measured by SWATH-MS, 97.9 % had corresponding genes in the gene expression datasets. P-values were adjusted for multiple hypothesis testing by Benjamini-Hochberg FDR correction; see Data file S4 for details. In this analysis we also validated INPP4B, CDK1 and ERBB2 (the proteins selected in the proteotype classification tree) as differentially regulated between ER^+^ vs. ER^−^ tumours, high vs. low grade tumours, and HER2^+^ vs. HER2^−^ tumours, respectively.

Second, we correlated log2 protein fold-changes (log2FC) as obtained from pairwise group comparison with the respective transcript log2FCs from the same comparisons in the five transcriptomic data sets. The same analysis was performed also for a subset of ERBB2, INPP4B and CDK1 protein-transcript pairs.

Third, a decision tree based on gene transcript levels was constructed in order to classify the samples into the five conventional subtypes and thus to compare the resulting model in terms of performance to the model (decision tree) based on proteotypes. In other words, we asked whether transcriptomics data are better at predicting the conventional subtypes. For this purpose, the same procedure as described above (“Construction of the decision tree” section) was applied on 1036 most variable (top 5%) probesets representing unique gene symbols and a set of 474 samples.

Fourth, preprocessed Level 3 RNA-seq data were downloaded from the TCGA data portal (https://portal.gdc.cancer.gov). Filtering and normalization was performed using edgeR package (Robinson et al., 2010; McCarthy et al., 2012). Limma (Ritchie et al., 2015) “RemoveBatchEffect” function was executed on log_2_ transformed Count Per Million (CPM) data. Batch corrected log_2_CPM values were then used in order to validate the hypotheses derived from our analysis of proteomic SWATH-MS data also on transcript level. A subset of 1078 samples with available information on gene expression, ER status and HER2 status were used to perform analysis of differential expression between the categories of clinical-pathological variable: 791 ER^+^ vs. 237 ER^−^ patients were compared in term of *INPP4B* expression, and 161 HER2^+^ vs. 554 HER2^−^ patients were compared in term of *ERBB2* expression using Wilcoxon rank sum test. As the information on tumour grade was unavailable for the dataset, we performed Spearman correlation of *CDK1* expression with expression data on commonly used proliferation marker *MKI67*.

##### h. Analysis of patient survival

Survival analysis was performed using Kaplan-Meier Plotter (http://kmplot.com) for relapse-free survival (RFS) involving a microarray dataset from 3951 breast cancer tissues (2018 database version) (Gyorffy et al., 2010). Each gene was represented by user-defined probe set, Affymetrix IDs were as follows: 205376_at (*INPP4B*), 203213_at (*CDK1*, referenced as *CDC2* in kmplot database), 216836_s_at (*ERBB2*), for reference genes 205225_at (*ESR1*) and 212023_s_at (*MKI67*). The population was split into high and low expression groups based on the incidence: (i) upper tertile for *INPP4B* and *ESR1* based on approximate proportion of ER^+^ and ER^−^ tumours, (ii) median for *CDK1* and *MKI67* genes based on approximate proportion of high and low grade tumours, and (iii) lower quartile for *ERBB2* based on approximate proportion of HER2^+^ and HER2^−^ tumours in the breast cancer population. 120 months follow up threshold was applied.

##### i. Statistical analysis of the IHC data

Associations between ERBB2 staining intensity and ER status were assessed by Fisher’s exact test in R 3.0.2.

## Supplemental information

Fig. S1. Overlap of cancer-related proteins identified by SWATH-MS and iTRAQ.

Fig. S2. Overview of samples from independent transcriptomics datasets DFHCC, DFHCC2, IRB, PNC and SUPERTAM_HGUPLUS_2 and how they were used for comparisons with the proteomic data and to build a decision tree.

Fig. S3. Decision tree based on gene expression data.

Fig. S4. Independent validation of INPP4B, CDK1 and ERBB2 association with ER status, tumour grade, and HER2 status (full version).

Fig. S5. Relapse-free survival (RFS) in breast cancer patients with high vs. low expression of the three key genes *INPP4B, CDK1, ERBB2*.

Fig. S6. Comparison of the number of proteins consistently quantified across samples by SWATH-MS and SuperSILAC.

Tab. S1. Coverage and abundance ranks of 19-protein signature identified by SuperSILAC compared to the abundance rank of the same proteins in our SWATH-MS dataset.

Data file S1. Sets of breast cancer tissues used in SWATH-MS study and for immunohistochemical validation.

Data file S2. Assay library for quantifying breast cancer associated proteins by SWATH-MS. Data file S3. Quantitative results of SWATH-MS study on 96 breast cancer tissues.

Data file S4. Comparison of protein and transcript levels.

Data file S5. Validation of SWATH-MS quantitation through S/MRM quantitation of 16 peptides representing 9 proteins in the same set of 96 breast tumors.

Data file S6. Manual validation of the three key proteins INPP4B, CDK1, and ERBB2.

